# Interactions between Submicron Carbon Particles, *Escherichia coli* and Humic acid with Plastic Surfaces

**DOI:** 10.64898/2026.02.26.708228

**Authors:** Nathan Bossa, Kobi Talma, Fiza Pir Dad, Lijia Gao, Gulsum Melike Urper-Bayram, Waqas ud Din Khan, Mark Wiesner

**Author notes:** **Corresponding Author:** Nathan Bossa [ ].

## Abstract

Plastic materials are widely used in engineered systems and increasingly accumulate in natural environments, where their surfaces interact with colloids, microorganisms, and dissolved organic matter. However, the relative roles of plastic surface properties versus particle-specific characteristics in governing organic matter retention remain poorly constrained. Here, attachment efficiency (α) was used to quantify intrinsic particle–collector affinity on three common thermoplastics (ABS, HDPE, HIPS) and glass beads as an inorganic reference. Surface chemistry, hydrophobicity, roughness, and charge were characterized, and interactions with submicron carbon particles (SCPs) and *Escherichia coli* were evaluated using column experiments. Extended DLVO (XDLVO) theory was applied to predict interaction energy barriers, and humic acid (HA) adsorption was quantified through batch isotherms.

XDLVO modeling predicted higher affinity of particles for plastics relative to glass; however, experimentally measured attachment efficiencies were uniformly low (α < 0.05) across all materials. Attachment was primarily governed by particle size and surface charge rather than collector hydrophobicity, roughness, or surface chemistry. SCP consistently exhibited higher α than bacteria, while differences among plastics were minor. Similarly, HA adsorption was weak and near-linear, with uptake following ABS ≈ HIPS > HDPE > glass, indicating reversible, partitioning-like association dominated by polymer-specific functionality rather than electrostatics. The absence of correlation between α and XDLVO-predicted energy barriers further demonstrates limitations of classical physicochemical models in describing particle– plastic interactions.

Collectively, these results indicate that pristine thermoplastic surfaces exhibit intrinsically low affinity for organic matter and that particle-specific properties dominate retention under low ionic strength conditions. Enhanced accumulation in environmental systems likely requires surface aging or conditioning processes not captured by classical interaction theory.

## 1. Introduction

Plastics are among the most pervasive synthetic materials in both natural and engineered environments. ^1,2^ Their durability, chemical resistance, and low cost have made polymers indispensable in product manufacturing and engineering systems. ^3^ However, these same properties also contribute to their environmental persistence, leading to the widespread accumulation of plastic debris in terrestrial and aquatic systems^4^. In engineered systems, such as water and wastewater treatment facilities, water distribution networks, and water storage infrastructure, plastics are commonly used in pipes, filters, and sensor housings where the interaction of plastic surfaces with suspended particles, bacteria, and dissolved organic matter can influence the performance of these systems ^5^. For example, bacterial attachment and biofilm formation on plastic pipes can deteriorate water quality and alter flow conditions, while adsorption of organic or colloidal particles affect membrane or sensor efficiency ^6^. In natural and agricultural systems, large quantities of mis-managed plastic materials are found at the macro-to nano-scales^7^. Plastic surfaces in the environment interact with colloids, organic matter, and microorganisms, altering contaminant transport, microbial colonization, and overall ecosystem function. For example, plastic is increasingly present in agricultural soils ^8^, both through intentional uses such as plastic mulches, irrigation materials, and coatings, and through unintentional contamination from plastic waste degradation and the accumulation of microplastics such natural settings, plastic surfaces.

Plastics comprise a highly diverse group of synthetic polymers differing in chemical composition, molecular structure, additives, and physical form, and they also exhibit a wide range of surface properties such as roughness, charge, hydrophobicity, and chemical reactivity. These surface characteristics strongly influence plastic aging ^9^, sorption of contaminants ^10^, microbial colonization, and interactions with environmental matrices across ecosystems.

Particle attachment and surface interaction processes are governed by a combination of physicochemical properties—including surface charge, hydrophobicity, and roughness—and solution conditions such as pH and ionic strength. While microbial adhesion to mineral or metallic surfaces has been extensively studied ^11^, comparable data for polymeric surfaces remain limited. Furthermore, few studies have systematically examined how both biological (e.g., bacteria) and abiotic carbon colloids (e.g., activated carbon) or dissolved organic matter interact with plastics under controlled conditions. Plastic surfaces are intrinsically heterogeneous, exhibiting microscale variations in topography, surface energy, and chemical functionality due to polymer crystallinity, additives, processing history, as well as heterogeneous surface environmental degradation (i.e. weathering, hydrolysis, dissolution, bio). Such heterogeneity induces spatially variable electrostatic potentials, hydrophobic domains, and nanoscale roughness that locally modulate the Derjaguin–Landau–Verwey– Overbeek (DLVO) interaction energy landscape ^12^. Accounting for this multiscale variability is essential to accurately predict particle–surface interactions and contaminant retention behavior on polymeric collectors. Understanding how the physicochemical properties and microscale heterogeneity of plastic surfaces affect the attachment efficiency of colloidal particles and the adsorption of ionic organic matter is essential for predicting interactions at the plastic interface and their subsequent behavior in environmental and industrial systems.

In this study, we investigate the attachment behavior of submicron carbon particles (SCP), *Escherichia coli* (*E. coli*), and humic acid (HA) on three representative plastics—ABS, HDPE, and HIPS—alongside glass beads (GB) as inorganic reference surface. The collectors were characterized using Fourier-transform infrared spectroscopy (FTIR), scanning electron microscopy (SEM), contact angle (CA) measurements, roughness measurements, and surface charge analysis to quantify their chemical and morphological properties. Furthermore, the interaction energy profiles were modeled using the extended Derjaguin–Landau–Verwey– Overbeek (XDLVO) theories. The attachment efficiency (α) of SCP and *E. coli* was quantified using column experiments, while humic acid adsorption was evaluated via batch studies. The attachment behavior of the SCP, *E. coli* and humic acid was compared with quantitative particle and surface properties, and qualitative analysis of surface energy obtained using XDLVO models.

## 2. Material and Methods

### 2.1. Collector Preparation and Properties

Pre-production plastic pellets, acrylonitrile butadiene styrene (ABS), high-density polyethylene (HDPE), and high-impact polystyrene (HIPS) were purchased from McMaster Carr Supply Company, IL, USA while, glass beads (GB) were purchased from VWR, USA. The plastic pellets were prepared for the attachment experiments by washing in 70% ethanol, followed by thoroughly washing with Milli-Q water and air drying in a laminar flow hood. The glass beads were prepared as described by Pelley and Tufenkji ^13^.

The surface morphologies of the collectors (ABS, HDPE, HIPS and GB) were examined by scanning electron microscope (SEM; Apreo S by ThermoFisher Scientific) at the Shared Materials Instrumentation Facility (SMIF), Duke University, USA. The accelerating voltage set at 2.00 kV. All samples were thoroughly dried and gold sputtered prior to SEM analysis and imaged at room temperature. Fourier transform infrared spectroscopy (FTIR) was used to characterize the chemical structure of samples. Measurements were performed using a ThermoFisher Nicolet iS50 FTIR spectrometer equipped with RaptIR+ accessory at SMIF Duke University, USA. Contact angle measurements were conducted using a contact angle tensiometer (Attention Theta Flex, Biolin Scientific, Duke University, USA). A droplet of deionized (DI) water (∼2µL) was pSCPed on the surface of each sample (ABS, HDPE, HIPS, and GB). The measurement range of the goniometer was 0-180°, with an angular deviation of (± 0.1°). Surface roughness was measured using a 3D optical/laser profilometer (VK-X3000, Keyence, SMIF Duke University, USA). Images approximately 210 µm x 280 µm were captured using a 50X objective. Material surface charge was measured using an Anton Paar SurPASS solid surface zeta potentiometer across a pH range from 2.7 to 7.8, and a Boltzmann curve was fit to the zeta potential data.

### 2.2. Colloid Preparation and Properties

Suspensions of carbon-based submicron particles (SCP) were used as an abiotic comparison with the microbial suspensions used in this work. The SCP (REGENESIS) is a commercial product used as an agricultural soil supplement. The materials is described by the manufacturer as Liquid PAC (powdered Activated Carbon), a bioremediation Products, formulated with 31–35% thermally activated, bituminous coal-based carbon for potable and wastewater tapplications.cations . The size of the SCP was measured on a Malvern Zetasizer Nano ZS (Malvern, U.K.) as described in Rogers et al. (https://dx.doi.org/10.1021/acs.est.3c03700). Measurements were taken at 25 °C using a reflective index of 1.50 for AC. The cell type used for this measurement was disposable folded capillary cells (DTS1070). Measurments were done in triplicate, with theconductivity, positioning, and duration automatically set. Zeta potential values were also measured using a Malvern Zetasizer Nano ZS (Malvern, U.K.). pH values were varied between 3 to 11 by adding 0.1M hydrochloric acid (HCl) and 0.1M sodium hydroxide buffer (NaOH).

The model bacterium used for this study was *Escherichia coli* K-12 (*E. Coli*), obtained from the Gunsch Lab at Duke University. A glycerol stock was stored at -80°C throughout the study. The model organism was grown in Lennox Luria Broth (LB) media to a stationary phase at 37°C with shaking. Upon reaching stationary phase, the liquid culture was centrifuged in 50mL tubes at 3000xg and 4°C for 10 minutes. Then, the LB media was aspirated from the tubes, and the pelleted bacteria was resuspended in EPA Moderately Hard water (table S.1). The size of *E. coli* was measured using a Malvern Mastersizer (Malvern, U.K.) and the zeta potential was determined using a Malvern Zetasizer Nano ZS (Malvern, U.K.), calculated from electrophoretic mobility measurements.

### 2.3. Column Experiments

A column setup, previously described by Rogers et al., ^14,15^was used to investigate particle attachment Briefly, the material to be tested was packed into a glass column (25mm x 150mm, Diba Omnifit EZ Chromotography Column) to the 11mm line creating a porous medium composed of either plastic or glass bead particles. Three plastic materials - high-density polyethylene (HDPE), acrylonitrile butadiene styrene (ABS), and high-impact polystyrene (HIPS) - along with glass beads (GB) were selected for this study. A syringe pump was used to pump DI water into the packed column from the bottom until the column was saturated.

#### Bacteria deposition

Bacteria suspensions monitored directly using an in-line ultraviolet-visible spectrophotometer at 600 nm (Evolution UV-Visible Spectrophotometer, ThermoFisher Scientific, USA). and a bypass to the column to measure the influent concentration, C_0_.. All bacteria from the tubing were removed by flushing with DI water. At least three pore volumes of background solution (EPA Moderately Hard water) were passed through the column to ensure the column was saturated with the background solution. The bacterial suspension was passed through the column with a 2.95mL/min flow rate, and the column effluent absorbance (at 600nm) was recorded every 10s for at least 4 pore volumes.

#### SCP deposition

1mL of the SCP feed suspension was collected to determine the influent concentration (C_0_). The SCP suspension was passed through the column at a flow rate of 1.5 mL/min, and the column effluent was collected in Falcon tubes using an automated sample collector, changing Falcon tubes every 45 seconds. The collected effluent was then pipetted into a 96-well plate and absorbance at 306 nm was measured using a microplate reader (SPECTROstar Nano, BMG Labtech, Germany).

### 2.4. Calculation of Attachment Efficiency (α)

Attachment efficiency, known as α, was used to evaluate the affinity of particles (SCP & *E. coli*) to the tested surfaces (ABS, HDPE, HIPS & GB). The attachment efficiency describes the likelihood that collisions between particles and collector surfaces result in attachment ^16^. The attachment efficiency is defined as:

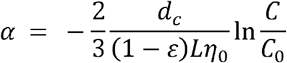

where *d*_*c*_ is the collector diameter, *L* is the column length, *ε* is the column porosity, *η*_0_ is the single collector contact efficiency, and 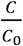 is the value of the plateau region in the breakthrough curves from the column experiments.

Using positively-charged aminated silica particles (nanoComposix, San Diego, CA) the same size as the test particle (1 µm for *E. coli* and 300 nm for SCP), and assuming that α = 1 for the positively-charged silica particles, or every collision results in attachment, the attachment efficiency (α) can be calculated using the following equation (table. S2):

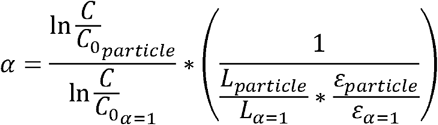

### 2.5. Extended DLVO Modelling

The classic Deraguin-Landau-Verwey-Overbeek (DVLO) theory models the interaction between two surfaces as the sum of van der Waals interactions, electrostatic interactions, and the Born repulsion energy (which is only influential at very small separation distances) ^17^. The extended DLVO (XDLVO) theory incorporates the polar interaction energies which occur in aqueous solutions, using the Lewis acid-base approach ^18^. The total interaction (in Joules) between surfaces over a separation distance, *h* (in meters), is described by ^19^:

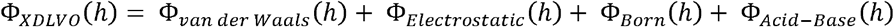

The equations used to calculate the components of the total interaction energy are presented in the Supporting Information. The interaction energy profiles were calculated using XDLVO theory, assuming that the interaction occurs between a spherical particle and a plate. This assumption is made given that the size of the SCP and *E. coli* particles is orders of magnitude smaller than the size of the plastic and glass collectors. The details of calculation is presented in SI.

### 2.6. Humic acid Solution Preparation and Adsorption

International Humic Substances Society Pahokee peat humic acid (PHA; Standard, 1S103H), was used as received without further purification. Stock solutions were prepared by dissolving 50 mg of humic substance powder in 2 mL of 0.1 N NaOH (ACS grade; Riedel-de Haen) and allowing the solution to equilibrate for 24⍰h prior to dilution. This procedure ensured complete dissolution and rehydration of the humic substance.

Working solutions for adsorption experiments were prepared by diluting aliquots of the stock solution with Milli-Q water (resistivity ≥ 18 MΩ•cm), followed by stirring for 1⍰h and standing for at least 24⍰h before use. The solution pH was adjusted using hydrochloric acid (HCl) or sodium hydroxide (NaOH) as required.

The kinetics of humic acid adsorption onto plastic pellets were evaluated using a humic acid solution with an initial concentration of 20 mg·L^−1^ at pH 6 and an ionic strength (I) of 0.05 M NaCl. Samples were collected at predetermined time intervals (0.25, 0.5, 1, 2, 5, 8, 12, 24) with a maximum contact time of 24 h. following established protocols ^20^

After 24 h of agitation, the equilibrium concentration of humic acid remaining in solution was determined by centrifugation at 7500 rpm for 10 min, followed by filtration through 0.45 μm syringe filters. The filtrates were analyzed using UV–visible spectroscopy (SPECTROstar Nano, BMG Labtech, Germany) at a wavelength of 310 nm, following previously reported methods ^21^.

Control samples containing humic acid solution without plastic were prepared at the same pH and ionic strength and analyzed using the same procedure. All calculations use the 24 h absorbance values to determine equilibrium concentration (C_e_) and adsorbed amount (q_t_), assuming linear UV-Vis response and the standard batch adsorption equation:

The amount of humic acid adsorbed on to plastic at time *t, q*_*t*_ (mg·g^−1^) was calculated as:

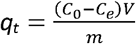

where *C*_*0*_ (mg·L^−1^) is the initial concentration of humic acid, *C*_*e*_ (mg·L^−1^) is the concentration at time *t* (h), *V* (L) is the solution volume, and *m* (g) is the mass of plastic.

Adsorption experiments were conducted at a plastic concentration of 100 g·L^−1^. All experiments were performed in triplicate at a constant temperature of 25 ± 1°C under continuous agitation at 250 rpm using a 211DS shaking incubator (Labnet International Inc., New Jersey, USA).

### 2.7. Statistical Analysis

Kruskal-Wallis test with Dunn’s multiple comparisons test was used to determine the dependence of bacterial and activated carbon attachment on material type. Two-way analysis of variance (ANOVA) with Bonferroni multiple comparisons test was used to evaluate the impact of particle type (Activated Carbon vs. Bacteria) on attachment to the same material. Correlation coefficients were calculated using Microsoft Excel Data Solver.

## 3. Results and Discussion

### 3.1. Material surface properties

The plastic materials used in this work, represent a range of plastic properties. HDPE is one of the most commodity plastics, valued for its mechanical strength, chemical resistance, and extensive use in bottles, pipes, films, and durable goods. ABS is a specialty engineering thermoplastic composed of acrylonitrile, butadiene, and styrene; its amorphous structure provides high toughness, rigidity, and chemical resistance, leading to widespread use in automotive components, electronics, appliances, and toys. HIPS is a niche styrenic polymer, consisting of polystyrene modified with rubbery butadiene domains to enhance impact resistance, and is commonly used in packaging, displays, and appliance interiors. Glass beads were used as a well-characterized reference medium as they provide a well-defined, reproducible silica surface that can be extrapolated to more heterogenous environmental materials such as sand and soil minerals. Fourier transform infrared spectroscopy (FTIR) analysis confirmed the manufacturer reported polymer compositions for all materials. ABS exhibited a characteristic nitrile (C⍰N) absorption band at ∼2230 cm^−1^ associated with the acrylonitrile component, along with aromatic C–H stretching (∼3020 cm^−1^) and aromatic C=C stretching (∼1600 and 1495 cm^−1^) from styrene, and butadiene-related bands near ∼965 cm^−1 22^. The FTIR spectrum of HDPE was dominated by aliphatic CH_2_ stretching vibrations at ∼2910 and 2850 cm^−1^, CH_2_ bending at ∼1460 cm^−1^, and a split CH_2_ rocking mode at ∼720 cm^−1^, indicative of crystalline chain packing ^23^. HIPS displayed aromatic C–H stretch (∼3020 cm^−1^) and C=C ring vibrations (∼1600 and 1495 cm^−1^) from the polystyrene backbone, along with butadiene-related bands such as C=C–H out-of-plane bending at ∼962 cm^−1^ and aromatic C–H out-of-plane bending in the range of ∼694–746 cm^−1^, characteristic of substituted benzene rings.^24^.

Surface hydrophobicity was assessed by static water contact angle measurements, yielding values of 111.20° ± 2.83°, 86.60° ± 4.53°, 118.92° ± 2.73° and 82.53° ± 2.59° for HDPE, ABS, HIPS and GB, respectively. HIPS exhibited the highest hydrophobicity, followed by HDPE, as both materials are primarily composed of nonpolar hydrocarbon groups (styrenic aromatic units in HIPS and –CH_2_– chains in HDPE), leading to low surface polarity and limited interaction with water. whereas ABS and GB were more wettable due to polar nitril group for ABS and silanol group for glass beads surfaces.

Despite their low to non-polar bulk chemistry, all plastic pellets exhibited low negative streaming potentials with values below -5 mV at pH 5.4 and 7.5, suggesting the presence of oxidize surface functionalities. At pH 5.4, the streaming potentials were –6.4, -7.9 and –7.0 mV for ABS, HDPE and HIPS, respectively. At ph. 7.5, these values decreased to – 8.6; - 13.2 and – 7.9 mV, respectively. The low negative surface charge is hypothesized to be the result of surface oxidation arising from thermal degradation during filament extrusion and palletization. High processing temperatures (typically 180–280 °C, depending on polymer) can induce C–H bond scission in the polymer backbone; in the presence of oxygen, free radical reactions lead to formation of oxidized surface groups such as carbonyl (–C=O), hydroxyl (–OH) and carboxyl (– COOH) functionalities ^25^. The typical peak of oxidation C=O ∼1710 cm^−1^, OH ∼3300 cm^−1^ were not detected on FTIR spectrum showing that oxidation certainly occurs only at the near plastic surface less than FTIR ATR typical penetration depth of 1 to 2 mm. The streaming potential values for GB were more negative than the three plastic surfaces, with values of –20.0 mV and –24.9 mV at pH 5.4 and 7.4 respectively.

Surface roughness was further quantified using optical profilometry at 50x magnification over a field of view of 208 × 278 µm. Out of the three plastic surfaces, HIPS presented smoothest surface with an average roughness (Ra) of 0.235 ± 0.05 µm. ABS was slightly rougher, but still relatively smooth with Ra of 0.305 ± 0.2 µm, while HDPE displayed a significantly higher roughness, almost 5–7× greater Ra than that of ABS and HIPS with roughness of 1.581 ± 0.7 µm. GB surfaces were smoother than the plastic surfaces, with a measured roughness of 0.108 ± 0.01 µm. Roughness measurements were performed after rigorous cleaning of the pellets using a 10% HNO_3_ acid solution followed by thorough rinsing to ensure roughness was not coming from adsorbed dust. Although absolute roughness values decreased for all materials after cleaning, the relative trend remained unchanged (Figure S.1), indicating that the elevated roughness of HDPE originates from intrinsic surface morphology rather than loosely adhered surface contaminants.

Scanning electron microscopy (SEM) images of the material surfaces are shown in Figure 1. ABS exhibited a predominantly flat surface decorated with small, rounded features up to ∼1 µm in size, along with numerous nanoscale markings (200 nm). HIPS surfaces presented heterogenous microscale features without a uniform pattern. In contrast, HDPE showed a pronounced valley-like morphology, characterized by flat ridge tops and narrow valley bottoms, consistent with its higher measured roughness. Glass beads exhibited largely smooth surfaces with occasional small surface defects.

**Figure 1.**
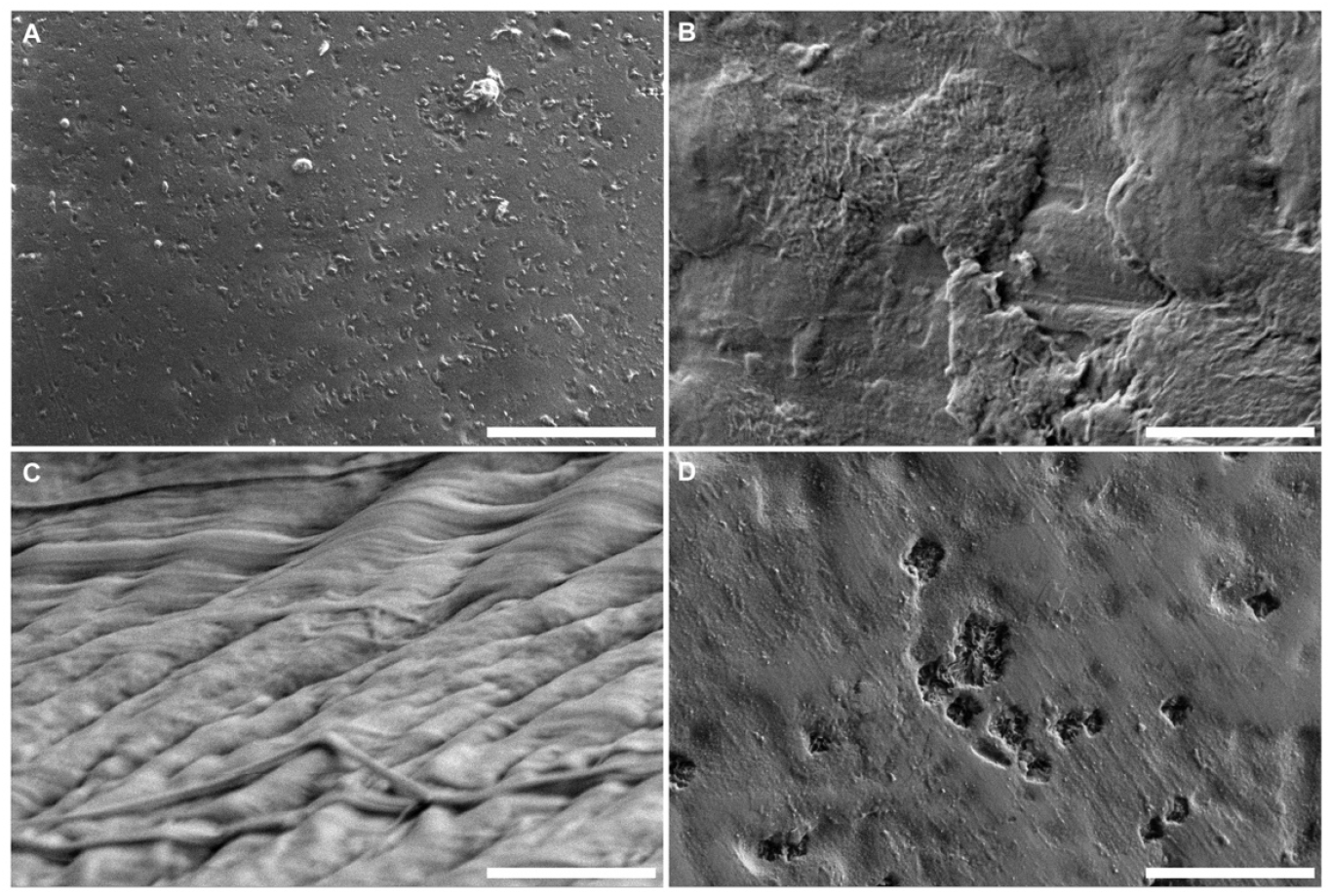
Scanning electron microscopy (SEM) images of plastic surfaces. Each frame corresponds to a different material: (A) acrylonitrile butadiene styrene (ABS), (B) high-impact polystyrene (HIPS), (C) high-density polyethylene (HDPE), and (D) glass beads (GB). All scale bars on graphs are 5 μm.

### 3.2. Particles properties

The submicron carbon particles (SCP), *E. coli* and humic acid were selected to represent a broad range of carbonaceous particle properties relevant to environmental and engineered systems. The SCP is a suspension of fine powdered carbon particles designed to adsorb contaminants such as organic compounds, metals, and per- and polyfluoroalkyl substances (PFAS). It is widely used in environmental engineering applications for groundwater and soil remediation^26^. *E. coli* was selected as a model microorganism due to its well-documented adhesion behavior and biofilm formation on diverse surfaces, providing insights into microbial interactions relevant to both environmental and engineered systems ^27–29^. While humic acid was selected because it represents a major fraction of natural organic matter (NOM), strongly influences contaminant mobility and microbial adhesion, and is commonly used as a model compound in environmental and water treatment studies ^30^.

The SCP suspension exhibited colloidal stability across a wide pH range. Its zeta potential remained consistently negative (–30 to –50 mV) between pH 3 and 11, with a value of –39.5 mV at its natural pH of 5.4. The average particle size was approximately ∼300 nm, with some particles extending into the micrometer range. Correspondingly, the low settling velocity (1.7 × 10^−6^ m/s) indicates that SCP particles remain well dispersed in suspension. The particles were irregular in shape with angular morphology (Figure S.2). *E. coli* exhibited a zeta potential of – 30.1 mV at pH 7.5, while the bacterial cells displayed a narrow particle size distribution centered at 1.15 µm (Figure S.3). Humic acid exhibited the expected negative surface charge and remained colloidally stable, with a hydration radius of approximately 150 nm due to electrostatic repulsion. The affinity of *E. coli* and SCP for the collector surfaces was evaluated using column experiments, whereas humic acid affinity was quantified using adsorption isotherm experiments. The characterization data was used to model the surface energy.

### 3.3. XDLVO Calculations of Particle-Surface Interactions

The extended DLVO (XDLVO) theory expands the classical DLVO framework by adding Lewis acid–base (polar) interactions—such as hydrogen bonding and hydrophobic effects—to the traditional van der Waals and electrostatic forces. This allows a more accurate description of particle–surface interactions in aqueous systems, where polar and hydration forces significantly influence adhesion and stability^18^. We use the XDLVO modeling for SCP and *E. coli* surface interaction but not for humic acid which, while it may be viewed as a nanoparticle, was treated here as an adsorbing solute.

XDLVO interaction energy profiles show the energy barrier to attachment, and detachment, as a peak known as the primary maximum (Φ_max_). The interaction energy profiles for SCP-collector and bacteria-collector interactions were calculated using XDLVO theory and the results are shown in Figure 2 and the values of Φ_max_ are shown in Table 1. For SCP particles, the predicted interaction energy barrier to attachment was highest for GB (69.63), while the interaction barrier to attachment was in a similar range for all three plastics (ABS = -0.43, HDPE = 7.52, and HIPS = 5.01). This trend was also seen for the *E. coli* particles, with interaction energy barrier for GB = 156.13, and similar range of values for ABS = -1.54 and HIPS = -1.00 while HDPE showed a mid-barrier value of 32.71. The results of XDLVO interaction energy barrier predict that attachment of both SCP and *E. coli* would be lower to GB as compared to the three plastic materials. The Φ_max_ values for the *E. coli*-collector interactions (between -1.54 and 156.13) are greater than those for the SCP-collector interactions (between -0.43 and 69.63), predicting that there would be less variation in attachment for SCP compared to *E. coli*. Based on the calculated XDLVO peak interaction energy values, it would be expected that SCP has greater attachment to the tested surfaces than *E. coli*, if attachment is driven solely by physicochemical properties.

**Table 1.** Calculated Φ_*max*_ values for sphere-plate models using XDLVO theory and the experimental conditions (SCP: pH = 5.4, I_s_ = 10^-3^ M; *E. coli:* pH = 7.5, I_s_ = 2.04×10^-3^ M)

| Material | SCP | *E. coli* |
| --- | --- | --- |
| | $\Phi_{\text{max}}$<br>($k_B T$) | $\Phi_{\text{max}}$<br>($k_B T$) |
| ABS | -0.43 | -1.54 |
| HDPE | 7.52 | 32.71 |
| HIPS | 5.01 | -1.00 |
| GB | 69.63 | 156.13 |

**Figure 2.**
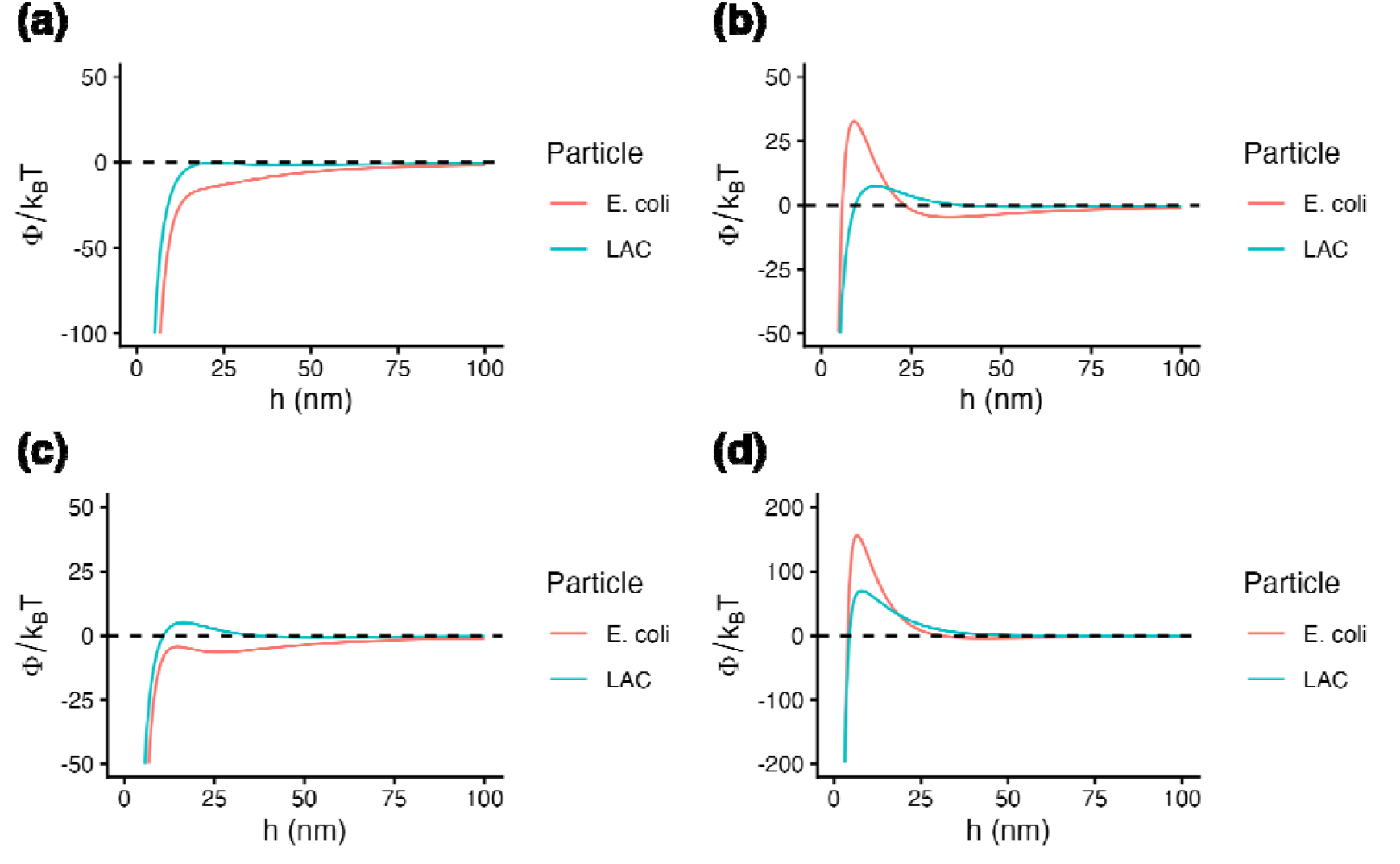
Predicted sphere to plate Φ_*XDLVO*_ interaction energy profiles for SCP and *E. coli* with (a) ABS, (b) HDPE, (c) HIPS, and (d) GB as a function of separation distance.

To evaluate the possibility of particle aggregation, the XDLVO interaction profiles for the case of sphere-sphere approximation were determined for SCP-SCP and *E. coli*-*E. coli* interactions and are shown in Figure S.4. The XDLVO theory clearly suggests that aggregation between like particles is not expected to occur under the experimental conditions.

### 3.4. Effect of material properties on particle attachment

At its core, α defines the probability of attachment following particle collision with another surface—the inherent affinity of SCP and *E. coli* for the tested surfaces (ABS, HDPE, HIPS, and GB). This parameter isolates the physicochemical interaction itself, independent of how often particles encounter surfaces. In contrast, the realized attachment rate for the column test is expressed as αη_0_L, where η_0_ captures the efficiency of a single collector in contacting particles flowing through the porous medium, and L reflects the length of the column. The parameters η_0_ and L can strongly amplify or dampen outcomes depending on particle size, mobility, hydrodynamics, and solution chemistry, but these factors do not alter the intrinsic stickiness encoded in α^31^.Thus, α is the fundamental driver of attachment, while η_0_L describe the physics of particle transport up to the surface of the porous medium “collector.” The product, αη_0_L— sometimes termed “relative attachment efficiency”—is therefore best viewed as a context-dependent expression of attachment. By contrast, α provides the universal descriptor of affinity, allowing one to separate the chemistry of attachment separate from the situational flow dynamics.^31^ Moreover, the parameter α allows for comparison with other studies of particle affinity for surfaces, including a rich literature on spherical polystyrene particle (now used as a surrogate for micro/nanoplastics) attachment to glass beads.

The results of α and C/C_0_ are presented in Figure 3 and Table 2. Detailed information of the calculation and the classic C/C_0_ curve is presented in the Supporting Information (Figure S.5). The intrinsic attachment probabilities (α) that value can range from 0 to 1, were consistently low for *E. coli* across the tested plastics, with values of 0.0185 for HIPS, 0.0181 for ABS, and 0.0154 for HDPE. These similar α values indicate that, despite differences in polymer type, the inherent affinity of bacterial cells for these surfaces is weak and does not differ substantially among them. However, the α value for GB was lower than that of the tested plastics (α = 0.00716). In contrast, liquid activated carbon exhibited higher α values, ranging from 0.0357 on HIPS, 0.0388 on HDPE and 0.0259 on ABS, up to 0.0506 on GB. These results show that activated carbon has a much greater intrinsic likelihood of attachment per collision compared to bacteria, with a particularly stronger affinity for glass beads. This finding supports the highest calculated XDLVO peak interaction energy values with SCP > *E. coli*. GB was predicted to present the lowest affinity, which was seen for *E. coli* but not for SCP. In addition, no significant difference was found for α between GB and other plastics. SCP and *E. coli*. Particle properties seem to be the main driver of α across the 4 materials while variation of collector properties do not seem to play a large role under these conditions.

**Table 2.** Calculated Values of α for Organic Particles to Various Collector Surfaces.

| Particle | Material | $C/C_0$ | $\alpha$ |
| --- | --- | --- | --- |
| SCP | ABS | $0.794 \pm 0.06$ | $0.0258 \pm 0.008$ |
| | HDPE | $0.717 \pm 0.03$ | $0.0388 \pm 0.005$ |
| | HIPS | $0.705 \pm 0.03$ | $0.0357 \pm 0.004$ |
| | GB<sup>*</sup> | $0.675 \pm 0.03$ | $0.0506 \pm 0.005$ |
| *E. coli* | ABS | $0.851 \pm 0.04$ | $0.0181 \pm 0.005$ |
| | HDPE | $0.877 \pm 0.03$ | $0.0154 \pm 0.004$ |
| | HIPS | $0.838 \pm 0.02$ | $0.0185 \pm 0.002$ |
| | GB<sup>*</sup> | $0.955 \pm 0.02$ | $0.00714 \pm 0.004$ |

**Figure 3.**
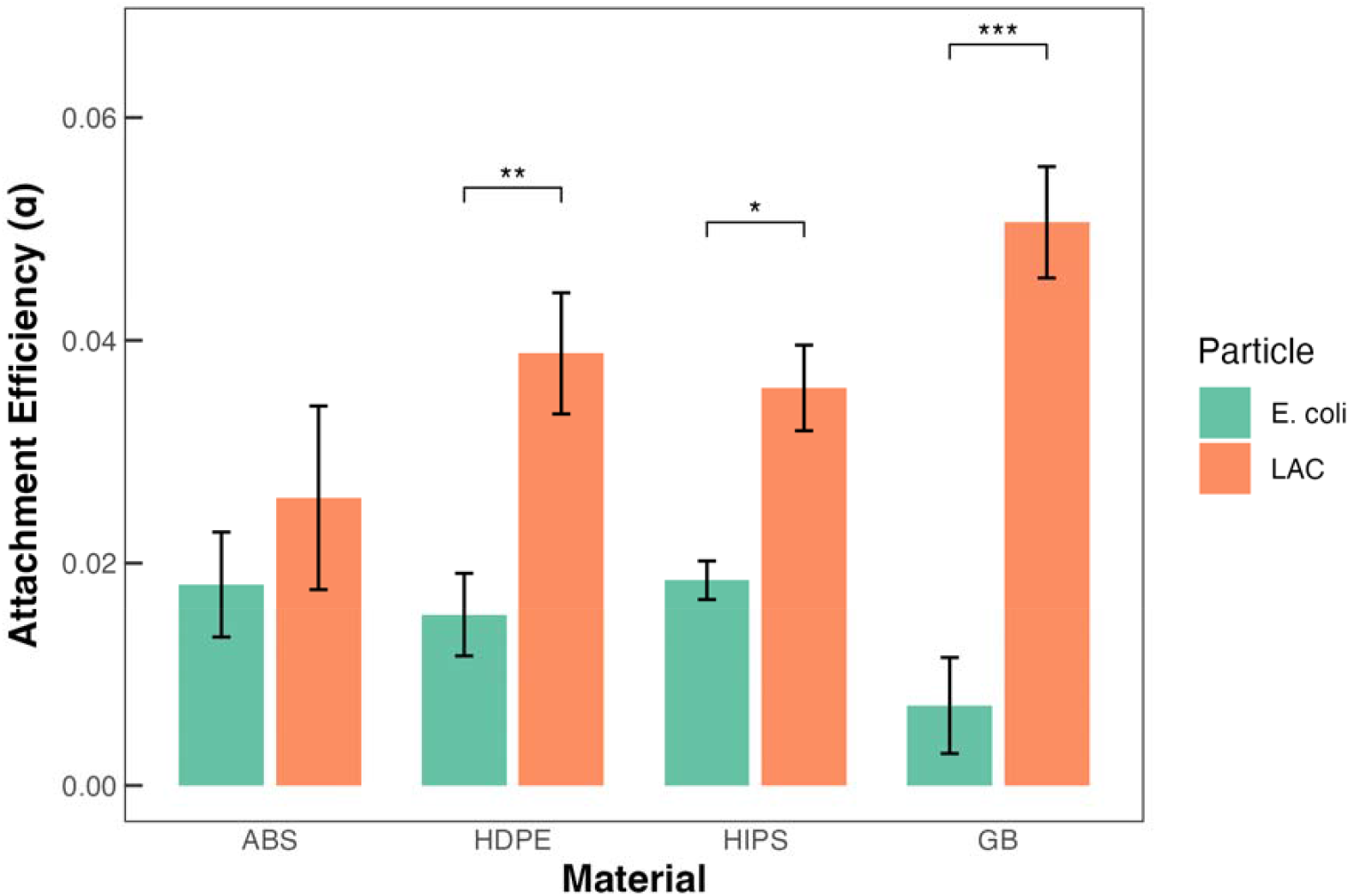
Measured attachment efficiencies for SCP and *E. coli* as a function of collector material.

Such α values are in the low range and represent a low affinity of *E. coli* and SCP toward plastic and glass surfaces. To our knowledge, α measurement of colloids toward plastic surfaces has been minimally investigated, although datasets on nano and microplastics transport in environmental porous media is available^32^. Attachment efficiencies (α) for plastic particles is found to range from 0.01 to 1.0, controlled mainly by ionic strength, particle shape, and organic coatings ^33,34^

Spherical polystyrene (PS) typically shows α = 0.01–0.04 at low salt concentrations (1 mM) and up to **0.8–1.0** at high salt (≥ 100 mM), while ellipsoidal PS reaches 0.3–1.0 due to enhanced interception and increased contact area^35,36^. Collector properties such as grain size, mineral composition, surface roughness, and coating strongly influence α: smooth quartz or glass beads promote lower attachment (α < 0.3), while rough or metal-oxide-coated sands enhance deposition (α approaching 1.0) by providing additional favorable surface sites and secondary energy minima^37^. Surface roughness and mineral heterogeneity create localized low-energy zones and secondary energy minima that favor attachment even under electrostatically unfavorable conditions^38^. In contrast, smooth, clean quartz or glass surfaces typically yield low α due to limited physical trapping and stronger electrostatic repulsion. Metal-oxide coatings (e.g., Fe or Al oxides) and organic matter on grain surfaces can either increase or decrease attachment depending on charge polarity and hydrophobicity. The difficulty remains the ability to measure and model such heterogeneity ^39^. We investigated the correlation of α with SCP and *E. coli* properties (i.e. particle size, particle charge), to collector properties (i.e. water contact angle, roughness and streaming potential) and finally the XDLVO energy barrier. Correlation analysis revealed that particle properties exerted stronger control over attachment efficiency (α) than collector surface properties under the tested conditions. Specifically, particle size and particle surface charge were strongly and negatively correlated with α (r = −0.860 and r = −0.855, respectively; p < 0.01), indicating that smaller and less negatively charged particles exhibited higher intrinsic attachment probabilities (Figure S6). In contrast, no statistically significant correlations were observed between α and collector characteristics such as surface roughness, contact angle, or streaming potential. Furthermore, α showed only a weak and statistically insignificant relationship with the XDLVO-predicted interaction energy barrier (Φmax), demonstrating that calculated physicochemical energy barriers did not quantitatively explain the observed attachment behavior (Table S3). These results reinforce that particle-specific properties dominate attachment under low ionic strength conditions and highlight limitations of classical XDLVO modeling in capturing the full complexity of particle–plastic interactions. (Figure S6).

### 3.5. Effect of materials properties on humic acid adsorption

Equilibrium adsorption isotherms of humic acid (HA) were determined for GB, ABS, HDPE, and HIPS after 24 hours of contact at five initial concentration levels (Figure 4). Adsorption parameters are summarized in Table 3.

**Table 3.**
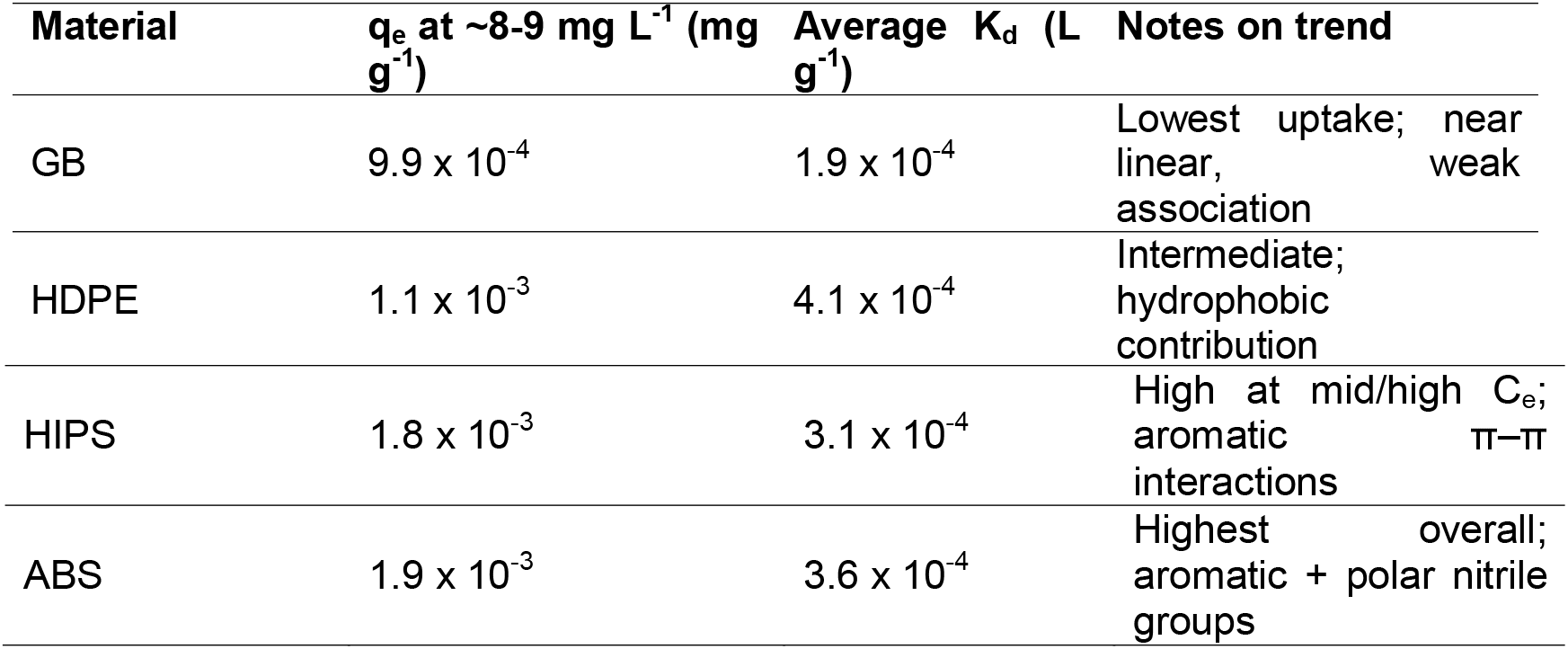
Equilibrium adsorption parameters of Humic acid on glass beads (GB) and plastic pellets (HDPE, ABS, HIPS) after 24 h contact time.

**Figure 4.**
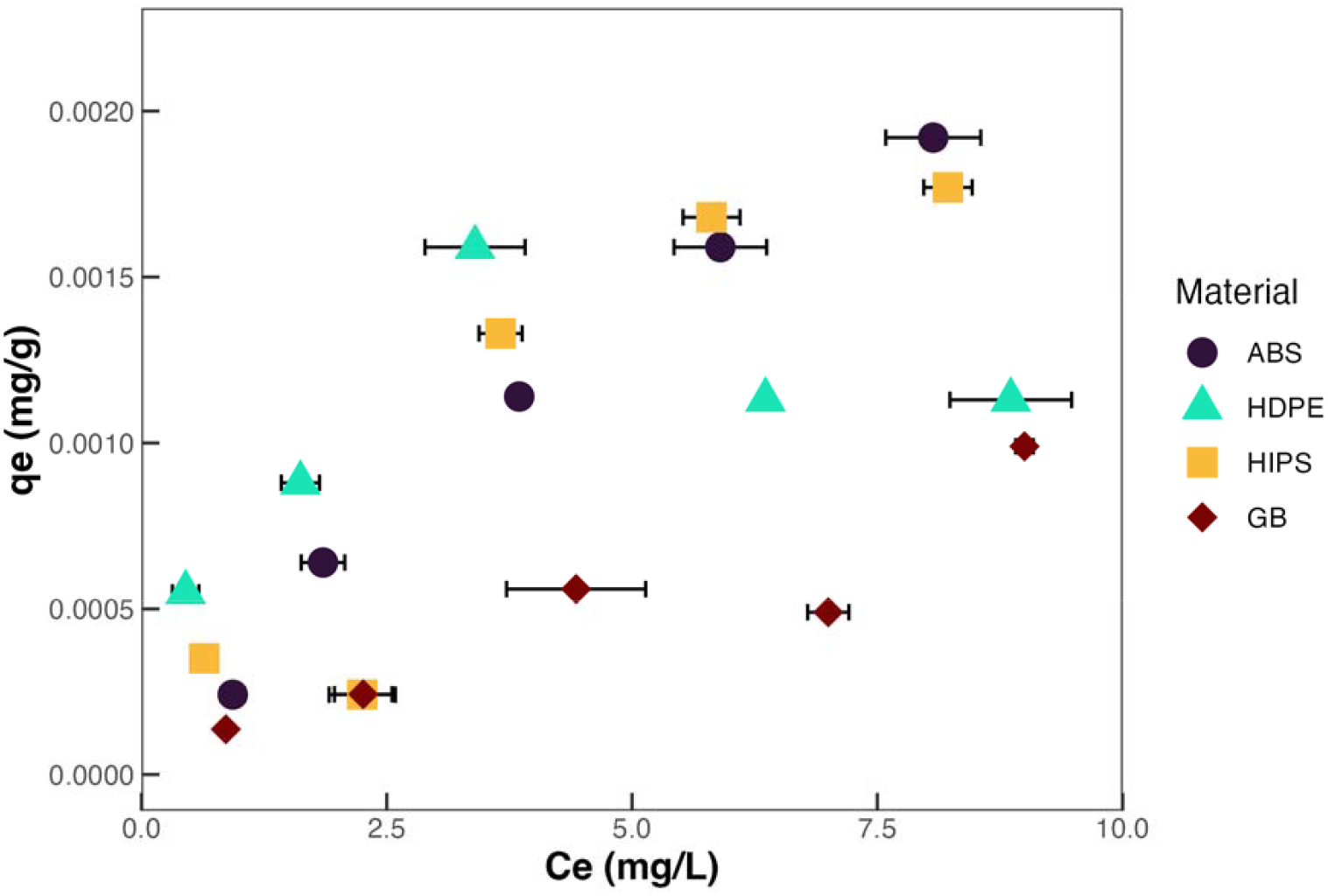
Equilibrium adsorption isotherms of Pahokee peat humic acid on glass beads (GB) and plastic pellets (ABS, HDPE, HIPS) after 24 h contact. Equilibrium adsorbed amount (q_e_) as a function of equilibrium aqueous concentration (Ce) at pH 6, I = 0.05 M NaCl, 100 g L^−1^ solid dose, 25 ± 1 °C. Symbols represent mean values (n=3); error bars denote standard deviation. Lines are guides to the eye. No adsorption plateau was observed within the studied range.

For all materials, the equilibrium adsorbed amount (q_e_) increased with increasing equilibrium aqueous concentration (C_e_), without reaching a plateau, over the investigated range (0.5-9 mg/L). At the highest equilibrium concentration (approximately 8–9 mg L^−1^), HA adsorption reached approximately 1.9×10−3 mg g^−1^ for ABS and 1.8×10−3mg g^−1^ for HIPS, compared to 1.1×10−3 mg g^−1^ for HDPE and 9.9×10−4 mg g^−1^ for GB. Apparent distribution coefficients (K_d_) confirmed enhanced affinity for polymeric surfaces relative to glass beads (ABS ≈ HIPS > HDPE > GB; Table 3). No adsorption plateau was observed, and isotherms exhibited approximately linear to mildly concave behavior for all materials within the investigated concentration range, consistent with apparent partitioning or weak, non-site-specific surface association rather than monolayer-limited adsorption.

Furthermore, short-term adsorption kinetics of humic acid (HA) were evaluated at an initial concentration of 5 mg L^−1^ over contact times ranging from 1 to 45 min. The lowest dissolved HA concentrations were observed between 10 and 15 min, with minimum remaining fractions of 68% for ABS, 75% for GB, 78% for HIPS, and 69% for HDPE. At longer contact times (30–45 min), the fraction of HA remaining in solution increased for all materials, reaching 77–79% for ABS, glass beads, and HIPS, and 87% for HDPE at 45 min (Figure S7).

(Average K_d_ = mean(q_e_/C_e_) across the full concentration range (∼0.5–9 mg L^−1^). q_e_ = equilibrium adsorbed amount (mg g^−1^), Values derived from UV-Vis absorbance at 310 nm after 24 h equilibration (pH 6, I = 0.05 M NaCl, 100 g L^−1^ solid dose, 25 ± 1 °C, n=3)

Short-term kinetic experiments revealed rapid initial humic acid (HA) uptake within the first 5– 10 min, followed by a partial increase in dissolved HA concentration at longer contact times. This rebound behavior was observed across all materials and was most pronounced for HDPE. The transient minimum in solution concentration suggests rapid initial association followed by partial desorption or redistribution of loosely bound HA fractions.

Taken together, the equilibrium and kinetic results indicate that HA association with pristine plastic and glass surfaces is weak and largely partitioning-like rather than site-limited. Across the concentration range investigated (0.5–9 mg L^−1^), adsorption increased approximately linearly with equilibrium concentration, with no evidence of saturation behavior. This near-linear response, combined with low uptake magnitudes (10^−3^ mg g^−1^), suggests reversible, non-specific interactions rather than strong chemisorption. The partial rebound in dissolved HA concentration during kinetic experiments further supports reversible association and dynamic redistribution among HA fractions. Apparent affinity followed the order ABS ≈ HIPS > HDPE > GB, indicating that polymer-specific chemical functionality exerts greater influence than hydrophobicity or surface charge alone ^40^. Enhanced uptake on ABS and HIPS likely reflects π–π interactions between aromatic styrenic domains and aromatic structures within HA, whereas HDPE—despite its high hydrophobicity—SCPks such specific interaction sites. The consistently lower adsorption on glass beads suggests that electrostatic repulsion limits HA association under these conditions. Together with the low attachment efficiencies measured for particulate organic matter, these findings indicate that pristine thermoplastic surfaces exhibit intrinsically limited affinity for both dissolved and colloidal organic matter.

## 4. Conclusion

This study demonstrates that unaged commercial plastic surfaces (ABS, HDPE, and HIPS) exhibit similarly low affinity toward diverse organic particles, including bacteria, liquid activated carbon, and humic acid. Attachment efficiency (α) measurements revealed that particle properties—particularly size and surface charge—dominate attachment behavior, while variations in plastic surface chemistry, hydrophobicity, and roughness play a secondary role under the tested conditions. Although XDLVO modeling predicted favorable interaction energies between organic particles and plastic surfaces, the experimentally measured attachment efficiencies (α) remained uniformly low across all materials. This discrepancy suggests that classic models FOR physicochemical interaction alone are insufficient to describe particle retention on polymeric collectors under flow conditions. XDLVO theory assumes chemically homogeneous, atomically smooth surfaces and does not account for microscale heterogeneity, nanoscale roughness, patchwise surface energy variation, or hydration and steric repulsion forces that may limit close approach. Moreover, even when primary minimum attachment is energetically favorable, hydrodynamic shear and torque within porous media can prevent stable adhesion or promote detachment of weakly bound particles. The relatively low ionic strength employed in this study further maintains electrostatic repulsion and reduces compression of the electrical double layer, limiting irreversible deposition. Consistent with these findings, humic acid adsorption experiments also demonstrated weak, near-linear association across all materials, with no evidence of site saturation and only modest differences between polymers and glass. The reversible and low-magnitude humic acid uptake further supports the conclusion that pristine plastic surfaces exhibit limited intrinsic affinity for organic matter under the tested conditions. Together, these factors likely explain why attachment remained low despite moderate hydrophobicity and weakly negative surface charge across plastics. These findings indicate that pristine thermoplastic surfaces are intrinsically resistant to organic particle accumulation and suggest that environmental aging, oxidative weathering, or biofilm conditioning layers may be necessary to substantially enhance particle retention in real-world systems. This work provides quantitative constraints for modeling plastic–organic interactions and underscores the need to incorporate surface heterogeneity and non-DLVO mechanisms in predictive frameworks.

## 5. Acknowledgements

This work was supported in part by the Engineering Research Centers Program of the National Science Foundation under NSF Cooperative Agreement No. EEC-2133504. This work was performed in part at the Duke University Shared Materials Instrumentation Facility (SMIF), a member of the North Carolina Research Triangle Nanotechnology Network (RTNN), which is supported by the National Science Foundation (award number ECCS-2025064) as part of the National Nanotechnology Coordinated Infrastructure (NNCI). The work and travel of authors was supported by the Higher Education Commission of Pakistan.

