## Supplementary material for "Interactions between Submicron Carbon Particles, *Escherichia coli* and Humic acid with Plastic Surfaces": Suplementary materials

Number of Pages: 9

Number of Figures: 7

Number of Tables: 3

**EPA Moderately Hard Water Preparation**

The following protocol was used to prepare 1 L of EPA Moderately Hard Water. 0.95 L of MILLI-Q, or equivalent, deionized water was placed in a properly cleaned carboy. 60 mg of MgSO_4_, 96mg NaHCO_3_, and 4mg KCl were added to the carboy and aerated overnight. 60mg CaSO_4_•2H_2_O was added to 0.05 L of MILLI-Q or equivalent deionized water in a separate flask. This flask was stirred on a magnetic stirrer until calcium sulfate was dissolved and added to the aerated 0.95 L and mixed well. The combined solution was aerated vigorously for an additional 24 hours to dissolve the added chemicals and stabilize the solution. Final prepared water quality parameters are shown in Table S.1.

**Table S.1:** Preparation of Synthetic Freshwater using Reagent Grade Chemicals^1^

|  | **Reagent Added (mg/L)^2^** | | | | **Approximate Final Water Quality** | | |
| --- | --- | --- | --- | --- | --- | --- | --- |
|  | NaHCO_3_ | CaSO_4_×2H_2_O | MgSO_4_ | KCl | pH^3^ | Hardness^4^ | Alkalinity^4^ |
| Moderately Hard | 96 | 60 | 60 | 4 | 7.4 -7.8 | 80 - 100 | 57 - 64 |

^1^Taken in part from Marking and Dawson (1973). ^2^Add reagent-grade chemicals to deionized water. ^3^Approximate equilibrium pH after 24 hours of aeration. ^4^Expressed as mg CaCO_3_/L.

**Calculation of XDLVO Interaction Energy Profiles**

The van der Waals interaction energy [J] can be calculated using the following expression:

$\Phi_{vdW}\left( h \right)= -\frac{Ar_{p}}{6h}\left[ 1+ \left( \frac{14h}{\lambda} \right) \right]^{-1}$

where *A* is the Hamaker constant [J], *r_p_* [m] is the radius of the spherical particle, and λ ≈ 10^-7^ m is the characteristic wavelength of the sphere-plate interactions. The Hamaker constant can be calculated using the minimum equilibrium distance between particle and material ($h_{0}$ = 0.25 nm) and the dispersive surface tension parameters of particle ($\gamma_{p}^{d}$), water ($\gamma_{w}^{d}$), and the material ($\gamma_{m}^{d}$):

$A=24\pi h_{0}^{2}\left( \sqrt{\gamma_{p}^{d}}-\sqrt{\gamma_{w}^{d}} \right)\left( \sqrt{\gamma_{m}^{d}}-\sqrt{\gamma_{w}^{d}} \right)$

The electrostatic interaction energy [J] between two spheres can be calculated as:

$\Phi_{EL}\left( h \right)= \pi\varepsilon_{r}\varepsilon_{0}r_{p}\left[ 2\Psi_{p1}\Psi_{p2}\ln\left( \frac{1+e^{-\kappa h}}{1-e^{-\kappa h}} \right)+\left( \Psi_{p1}^{2}+\Psi_{p2}^{2} \right)\ln\left( 1-e^{-2\kappa h} \right) \right]$

where $\varepsilon_{r}$ is the dimensionless relative dielectric constant of the suspending liquid, $\varepsilon$ [C^2^/(J*m)] is the vacuum permittivity, $\Psi_{p1}$ [V] is the surface potential of the spherical particle, $\Psi_{p2}$ [V] is the surface potential of the plate, and κ [1/m] is the inverse of the diffuse layer thickness:

$\kappa= \left[ \frac{2I_{s}N_{A}1000e^{2}}{\varepsilon_{r}\varepsilon_{0}k_{B}T} \right]^{\frac{1}{2}}$

where *I_s_* [mol/L] is the ionic strength, *N_A_* = 6.02 x 10^23^ [1/mol] is Avogadro’s number, *e* = 1.602 x 10^-19^ [C] is the elementary charge, *k_B_ =* 1.38 x 10^-23^ [J/K] is the Boltzmann constant, and *T* [K] is the fluid temperature.

The Born repulsion energy [J] for sphere-sphere interactions can be estimated by:

$\Phi_{Born}\left( h \right)=\frac{A\sigma_{Born}^{6}}{7560}\left[ \frac{8r_{p}+h}{\left( 2r_{p}+h \right)^{7}}+ \frac{6r_{p}-h}{h^{7}} \right]$

where $\sigma_{Born}$ [m] is the Born collision parameter, with 5 x 10^-10^m a commonly used value.

The Lewis acid-base interaction energy [J] can be estimated by:

$\Phi_{AB}\left( h \right)=2\pi r_{p}\lambda_{AB}\Phi_{AB(h=h_{0})}exp\left[ \frac{h_{0}-h}{\lambda_{AB}} \right]$

where $\Phi_{AB(h=h_{0})}$ [J/m^2^] is the Lewis acid-base free energy of interaction between two surfaces at “contact”, $\lambda_{AB}$ [m] is the decay length of water. $\Phi_{AB(h=h_{0})}$ can be estimated using the van Oss-Chaudhury-Good (vOCG) method:

$\Phi_{AB(h=h_{0})}=-2\left( \sqrt{\gamma_{1}^{+}\gamma_{2}^{-}}+\sqrt{\gamma_{1}^{-}\gamma_{2}^{+}} \right)$

where $\gamma_{1}^{+}$ and $\gamma_{2}^{+}$ [J/m^2^] are the electron-acceptor, or acidic, components of the surface free energies of surfaces 1 and 2, and $\gamma_{1}^{-}$ and $\gamma_{2}^{-}$ [J/m^2^] are the electron-donor, or basic, components of the surface free energies of surfaces 1 and 2.

**Additional Material Characterization**

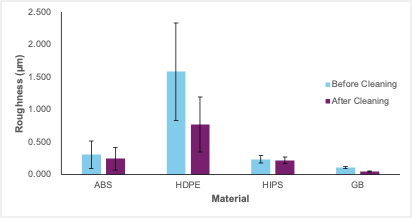

**Figure S.1:** Comparison of roughness measurements before and after cleaning using 10% HNO_3_, followed by thorough rinsing in Milli-Q water. Error bars show standard deviation of roughness measurements.

**Particle Properties Characterization**

All images/graphs of particle characterization that we don’t include in the main text

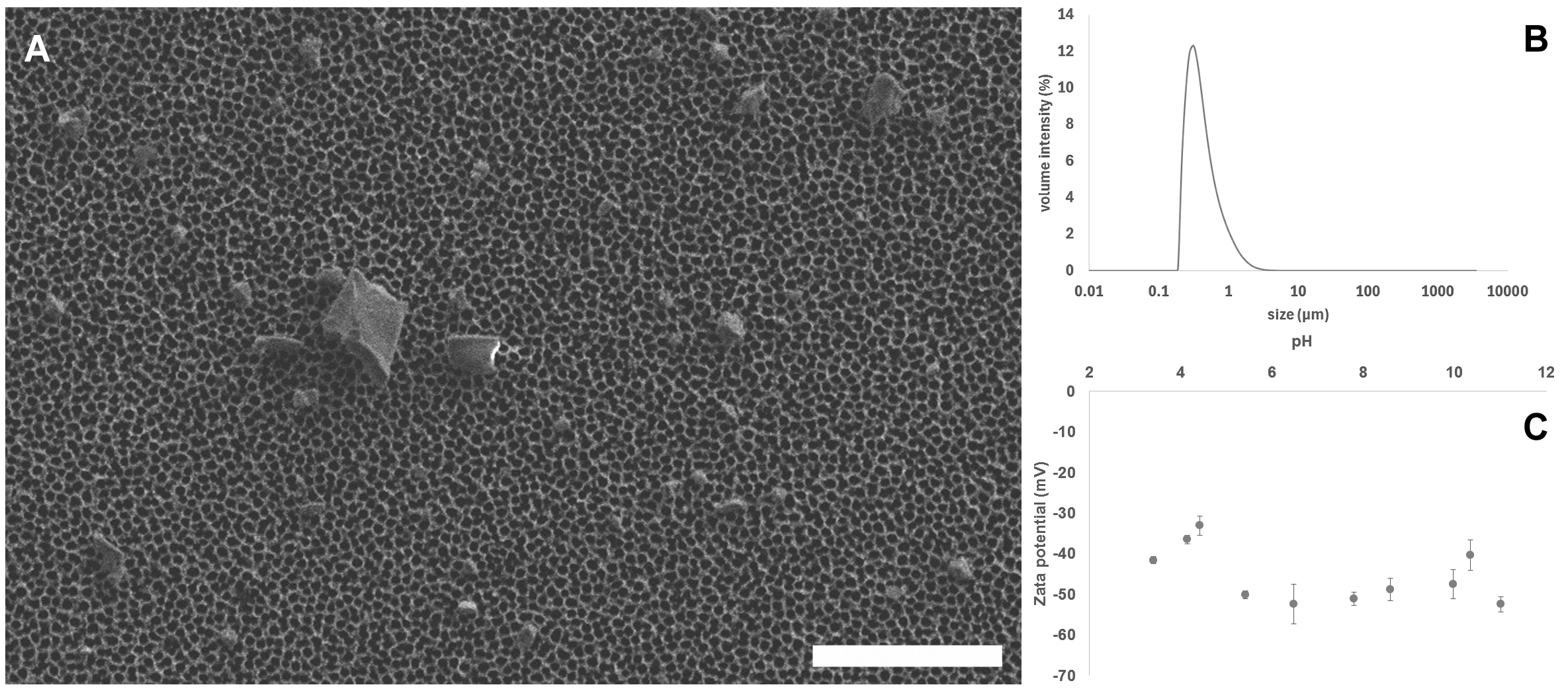

**Figure S.2:** Characterization of liquid activated carbon (LAC). (A) Scanning electron microscopy (SEM) image of LAC - scale bar 5 μm. (B) Dynamic light scattering (DLS) size distribution of LAC particles. (C) Zeta potential vs. pH of LAC.

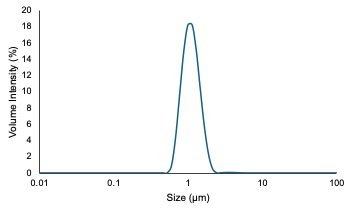

**Figure S.3:** Dynamic light scattering (DLS) size distribution of *E. coli*.

**XDLVO Interaction for Aggregation (sphere to sphere equations + interaction graphs)**

In order to evaluate the possibility of particle aggregation, the Φ_XDLVO_ interaction energy profiles for the sphere-sphere approximation as applied to identical bacteria-bacteria and activated carbon-activated carbon interactions were constructed. The XDLVO theory suggests that no aggregation between like particles is expected to occur under the experimental conditions.

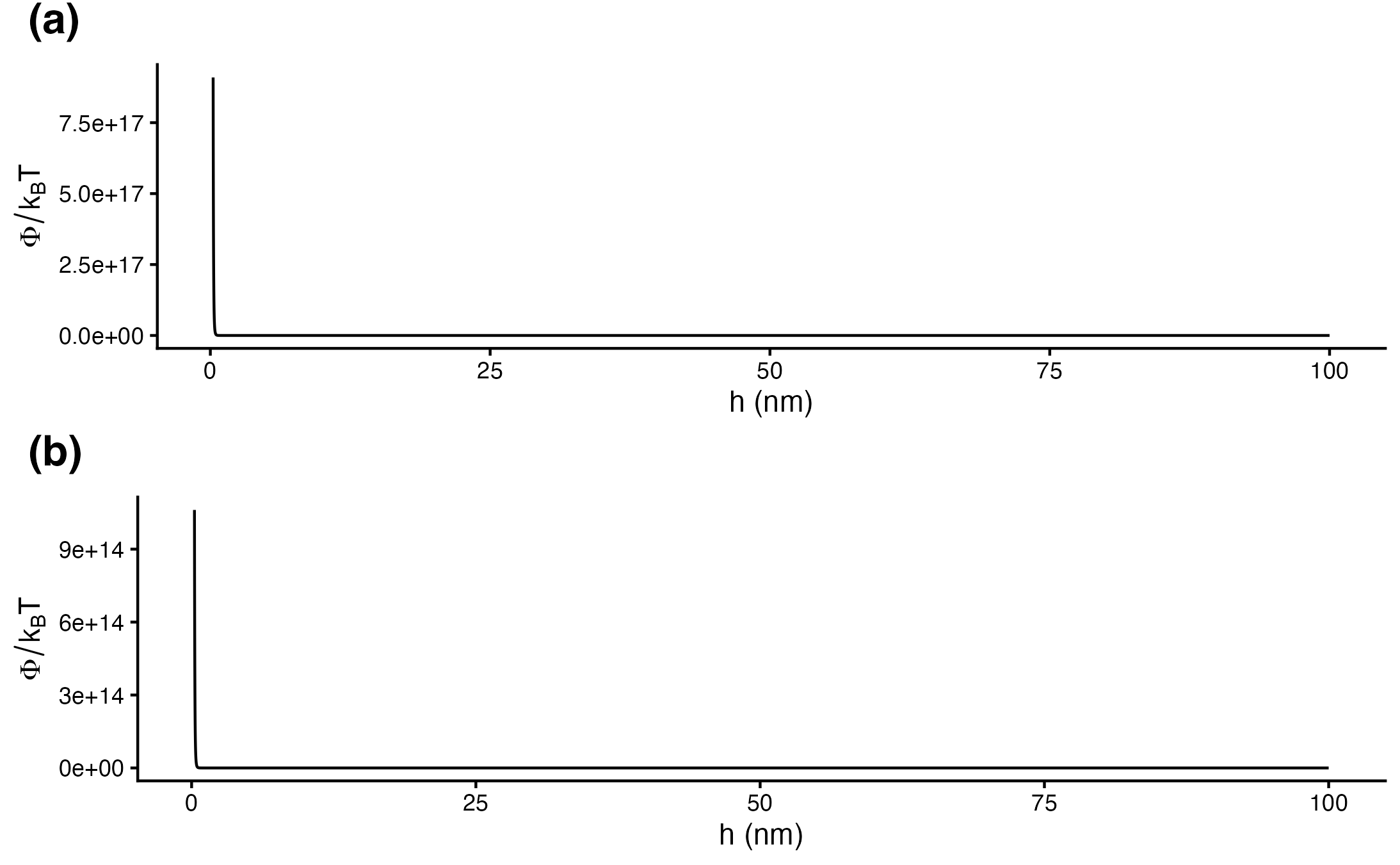

**Figure S.4:** Predicted sphere-sphere Φ_XDLVO_ interaction energy profiles for (a) *E. coli-E. coli*, (b)LAC-LAC as a function of separation distance, for the experimental conditions.

**Attachment Efficiency from Breakthrough Curve**

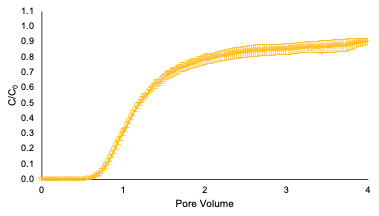

**Figure S.5:** Breakthrough curve for *E. coli* interacting with ABS. The error bars are standard error for triplicate measurements.

Positive particle columns were carried out using aminated silica particles (nanoComposix; San Diego, California) in order to calculate the attachment efficiency (α) using the normalization equation described in the main manuscript text. The positive particle experiments were performed as described in the Methods section with the following differences: (1) the column was packed to the 5cm line with material, (2) DI water was used as the background solution, and (3) absorbance was measured at 350nm. The C/C_0_ values reported in Table S.2 show that these particles have a high affinity to the test materials.

**Table S.2:** Measured Steady State Values from Positive Particle Breakthrough Curves

| **Material** | **C/C_0_** |
| --- | --- |
| ABS | 0.00800 ± 0.00352 |
| HDPE | 0.0115 ± 0.0124 |
| HIPS | 0.00929 ± 0.00110 |
| GB | 0.0132 ± 0.0145 |

**Correlation Results and Significance Determination**

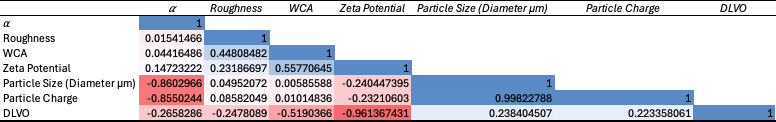

**Figure S.6:** Results of Correlation Analysis

**Table S.3:** Significance Evaluation of Correlation Analysis

| **Parameter** | **Correlation Coefficient** | **t – Statistic** | **p-value** |
| --- | --- | --- | --- |
| Roughness | 0.01541466 | 3.78E-02 | 0.9711021 |
| Water Contact Angle | 0.04416486 | 1.08E-01 | 0.9172985 |
| Surface Zeta Potential | 0.14723222 | 3.65E-01 | 0.7279031 |
| Particle Size | -0.8602966 | -4.13E+00 | 0.0061222 |
| Particle Charge | -0.8550244 | -4.04E+00 | 0.0068134 |
| Φ_XDLVO - Max_ | -0.2658286 | -6.75E-01 | 0.5245545 |

**Humic Acid Adsorption**

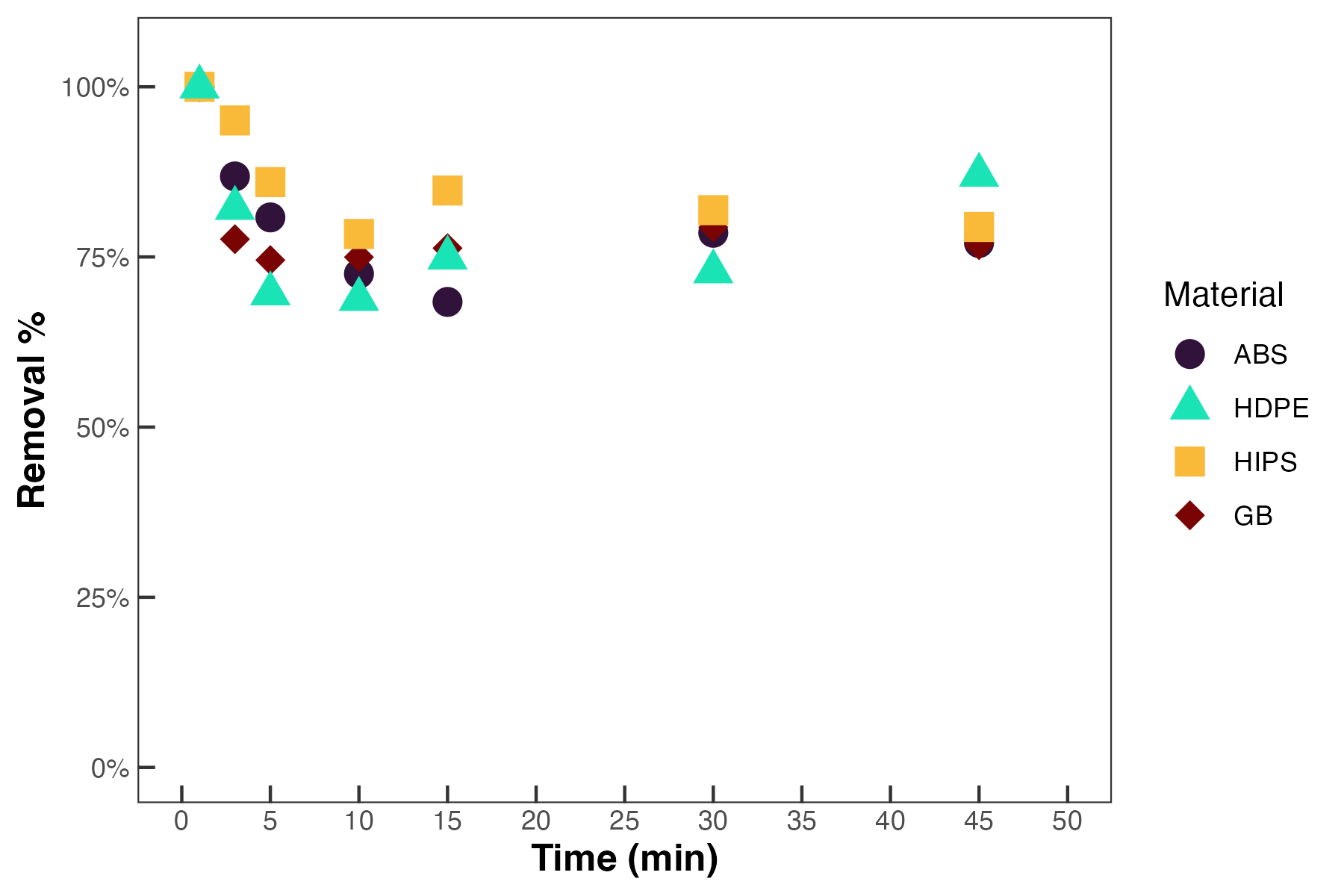

**Figure S.7:** Kinetic humic acid adsorption experiment results. Initial HA concentration = 5 mg L^-1^.
